# Species-specific responses of diurnal birds to nocturnal conspecific song playback

**DOI:** 10.64898/2026.08.27.747485

**Authors:** Kinga Buda, Jakub Buda, Michał Budka

## Abstract

The vast majority of birds are diurnal and concentrate their vocal activity during daylight hours. However, some diurnal birds can also be vocally active at night, although the functions of this phenomenon remain poorly understood. We conducted playback experiments in the Warta Landscape Park (central Poland) to determine whether nocturnal singing by diurnal birds serves breeding-related functions by analysing responses to playback of songs from unfamiliar conspecific males. The sedge warbler (*Acrocephalus schoenobaenus*) was selected as the focal species because it exhibits relatively high levels of nocturnal song activity, while nine additional diurnal species detected near focal sedge warbler territories were included to explore whether responsiveness to nocturnal conspecific song extended across a broader taxonomic range. Playback experiments were conducted during the early and late stages of the breeding season and during the early and late parts of the nautical night. Out of 10 species tested, three responded vocally: sedge warbler, Savi’s warbler (*Locustella luscinioides*), and common snipe (*Gallinago gallinago*). Sedge warblers did not modify song rate and song duration but increased flight activity after nocturnal playback. Savi’s warblers and common snipes produced more vocalisations after playback than before in May, a pattern consistent with territorial defence function. General nocturnal vocal activity was higher at the beginning of the season, suggesting that birds’ motivation to establish territories and form pairs can extend into the night, providing additional benefits. Moreover, the probability of singing by the sedge warbler was higher in the latter part of the night. Our study demonstrates that nocturnal stimulation of foreign male playback of diurnal birds can elicit vocal responses from conspecifics, suggesting that nocturnal singing can occur in the absence of obvious artificial light pollution, but in the case of some species and environmental conditions, it may contribute to nocturnal social communication, especially in the early breeding season.

## INTRODUCTION

Singing activity in many bird species is strongly structured over the daily cycle and often peaks around dawn, with some species also showing elevated activity around dusk (Catchpole & Slater, 2003). Several non-mutually exclusive explanations have been proposed for the dawn peak, including atmospheric conditions favourable for acoustic communication and limited foraging opportunities under low light levels, and increased opportunities for social signalling (Gil & Llusia, 2020; Kacelnik & Krebs, 1983). Although nocturnal vocal activity is commonly associated with nocturnal and crepuscular taxa, such as owls and nightjars, a growing body of evidence shows that many otherwise diurnal species also vocalise at night (Buda et al., 2025; Celis-Murillo et al., 2016a; La, 2012). Nocturnal song may serve social functions similar to those of daytime song, including mate attraction and territorial defence, although direct evidence remains limited to relatively few species (Dickerson et al., 2023). However, extending singing activity into the night may also entail costs. For example, higher nocturnal song rates in male common nightingales were associated with a greater overnight loss of body mass, indicating increased use of energy reserves (Thomas, 2002). Nocturnal singing may also increase a signaller’s exposure to nocturnally active predators: in an experimental study, predators, including owls and mustelids, approached nocturnal song playback, whereas no such approaches were recorded during daytime trials (Buda et al., 2024). Together, these potential benefits and costs suggest that the expression and function of nocturnal singing are likely to depend on species-specific and environmental contexts.

The reported prevalence of nocturnal singing by diurnal birds varies geographically. In Central Europe, 46 of 128 recorded bird species were detected singing at night; 35 of these 46 nocturnal singers (76%) were classified as diurnal (Buda et al., 2025). A review of 749 North American breeding bird species found reports of nocturnal vocalisations for at least 30% of species, more than 70% of which were classified as diurnal (La, 2012). By contrast, only three of 50 diurnal species sang during astronomical night in the Afrotropical highlands of Cameroon, producing a total of 10 songs (Budka et al., 2021). This may indicate that tropical birds engage in potentially risky behaviours less often than temperate species, possibly because of differences in life-history strategies or to reduce competition for acoustic space with other nocturnal animals, such as insects and anurans (Ghalambor & Martin, 2001).

Although nocturnal song may serve social functions similar to those of diurnal song (Catchpole & Slater, 2003; Dickerson et al., 2023), its role in otherwise diurnal birds remains poorly understood. By extending singing activity into the night, males may increase their total song output over a 24-h period and, consequently, their overall signalling effort. This may increase signal exposure to both potential mates and rivals, although it remains unclear whether nocturnal song output reliably indicates male quality. Two broad, non-mutually exclusive functional hypotheses follow from these potential receiver effects. Under the intersexual hypothesis, males may sing at night to advertise their presence or availability to prospecting females, including potential extrapair mates. Under the intrasexual hypothesis, nocturnal song may signal territory occupancy or mediate interactions with rivals. Evidence for the relative importance of these functions remains species-specific. In field sparrows (*Spizella pusilla*), males did not sing or countersing in response to simulated nocturnal intrusions, whereas females showed movement responses that varied across the breeding cycle, suggesting that nocturnal song may primarily serve an intersexual function in this species (Celis-Murillo et al., 2016b). In other species, nocturnal song may contribute to both mate attraction and territorial defence (Dickerson et al., 2023). The occurrence and intensity of nocturnal singing also vary across the breeding season and with environmental conditions, including weather and lunar illumination (Foote et al., 2017). Controlled playback experiments across a broader range of species and social contexts are therefore needed to determine whether and how diurnal birds respond to conspecific song at night.

In this study, we test whether and how otherwise diurnal birds respond to conspecific song broadcast at night using automated playback experiments. The sedge warbler (*Acrocephalus schoenobaenus*) was selected as the focal species because it exhibits relatively high levels of nocturnal song activity (Buda et al., 2025). We additionally included nine diurnal species detected near focal sedge warbler territories to explore whether responsiveness to nocturnal conspecific song extended across a broader taxonomic range. We hypothesised that, if nocturnal conspecific song functions as a socially relevant signal, birds would increase their vocal or movement activity after playback relative to the before-playback period. We further predicted stronger responses during the early- than the later-season sampling period, when territorial establishment and mate acquisition were expected to be more prevalent. Such seasonal variation would be consistent with a role of nocturnal song in territorial or mating interactions. We also examined whether nocturnal activity and playback responses differed between the early and late parts of the night. Finally, at sedge warbler territories, we recorded bird movements near the playback source as a complementary measure of behavioural responsiveness. A rapid vocal response, countersinging, or directed movement towards the playback source would be consistent with an intrasexual function involving male–male competition. The absence of detected aggressive responses would support an intersexual function of nocturnal singing, related to females attraction.

## METHODS

### Study site

The study was conducted in the Warta Landscape Park in central Poland. This conservation area is covered by forests, wet and semi-natural meadows, grasslands, reedbeds, oxbow lakes, riparian woodlands, wetlands and other habitats shaped by periodic flooding of the Warta River. Approximately 150 bird species breed in the park. The 25 experimental sites were established within areas occupied by singing male sedge warblers, in wetland vegetation adjacent to reedbeds. All experimental sites were located outside of the villages to reduce exposure to anthropogenic noise and artificial light pollution.

### Field data

Playback trials were conducted at the same experimental sites during two sampling periods, corresponding to the early (from 29 April to 3 May 2024) and late (from 4 to 14 June 2024) stages of the breeding season. Because birds were not individually marked, repeated trials at a site cannot necessarily be attributed to the same individual. All trials were conducted during nautical night – the period from nautical dusk to nautical dawn, when the solar centre was at least 12° below the horizon.

Experimental sites were established within sedge warbler territories because this was the focal species of the study. To identify additional species for playback trials, we conducted a 10-minute point count at each experimental site between sunrise and 10:00 (local time) from 27 to 28 April 2024. During each count, we recorded all bird species detected visually or acoustically within a 100-m radius. In addition to the sedge warbler, we selected three diurnal species at each site that were locally present and showed the highest vocal activity during the point count. This approach allowed us to use the same experimental locations and equipment to conduct an exploratory assessment of nocturnal responsiveness across a broader range of locally co-occurring species. In total, we selected and further tested 10 bird species. The number of sites at which each species was tested was: 25 sites for sedge warblers, 22 for Eurasian skylarks (*Alauda arvensis*), 16 for common snipes (*Gallinago gallinago*), 11 for Savi’s warblers (*Locustella luscinioides*), 9 for yellowhammers (*Emberiza citrinella*), 7 for common reed buntings (*Emberiza schoeniclus*), 3 for eurasian blackcaps (*Sylvia atricapilla*), 3 for willow warblers (*Phylloscopus trochilus*), 2 for common blackbirds (*Turdus merula*), and 2 for common whitethroats (*Curruca communis*).

Experimental sites were separated by at least 150 m. Sites tested on the same night were separated by more than 350 m to reduce the likelihood that birds at one site were stimulated by broadcast at another site. At each site, an Ultimate Ears Boom 2 loudspeaker connected to an Olympus LS–P1 player was mounted on a pole approximately 1 m above ground at the reedbed edge, within the area occupied by the focal sedge warbler male. An infrared camera trap (Camera Trap Full HD 40 IR + GPRS MMS) was mounted on a separate pole 3 m from the loudspeaker and 1.5 m above the ground, with its field of view centred on the area surrounding the speaker. The camera was used to record visible bird movements during the experiment. Additionally, vocal responses of the birds at each experimental site were recorded using two AudioMoth 1.1.0 autonomous acoustic recorders (mono wav files, 48 kHz/16–bit sampling rate), attached to vegetation 2–5 m from the loudspeaker and 1–2 m above ground.

All devices were programmed to operate automatically, and no observers were present at the experimental sites during trials, thereby minimising potential observer effects on bird behaviour. Each site was exposed to the same playback sequence twice per night. In May, early-night blocks were conducted between 22:30 and 00:00, and late-night blocks between 02:00 and 03:30. In June, the corresponding blocks were conducted between 23:30 and 01:00, and between 01:00 and 02:30, respectively. The Moon was above the horizon during part of the late-night blocks on the first three sampling nights (29 April–1 May); during all other trials, it was below the horizon. Trials were restricted to nights without precipitation and with low wind and cloud cover.

### Playback design

For each species, we selected five high–quality recordings from the Xeno-Canto sound archive. Recordings were selected according to the following criteria: high signal-to-noise ratio, minimal overlap with other species, absence of anthropogenic noise, known recording location, and geographical separation from the study population. Each recording represented a different individual. We preserved the natural temporal structure of the songs. When recording was shorthorn than 10 minutes, it was looped as necessary to create a 10-min species-specific playback sequence.

A site-specific playback block was prepared for each experimental site and comprised 10-min sequences for the sedge warbler and three additional species selected at that site. The four species-specific sequences were separated by 10-min silence periods during which no stimulus was broadcast, resulting in a total block duration of 90 min. The order of species within each block was randomised. The same block was broadcast twice per night during the early- and late-night periods. For each target species, the experimental trial comprised three phases: 10 minutes before the playback period, 10 minutes of playback, and 10 minutes after the playback period. For consecutive species in a block, the after-playback period of one trial also served as the before-playback period of the next trial.

All playback files were prepared by one person (KB) using Avisoft SASLab Pro 5.2.12 software (Avisoft Bioacoustics e.K., Glienicke, Germany). A high–pass filter was applied to reduce low-frequency background noise. The cutoff frequency was below the minimum song frequency of every target species. Playback amplitude was calibrated to 85 +/– 2 dB SPL(A) at 1 m from the speaker using a UNI–T UT351 sound-level meter. This amplitude was used as a standardised broadcast level across species.

### Acoustic and video analyses

Vocal responses recorded by autonomous sound recorders were analysed using Raven Pro 1.6.5 software (Cornell Lab of Ornithology, Ithaca, NY, USA). Spectrograms were generated using a 1024-sample Hamming window, with brightness and contrast set to 60%. Each song and call recorded during the experiment was identified to species during manual scanning of spectrograms and auditory verification of recordings. Only vocalisations of the target species were included in the species-specific analyses. We used recordings from one randomly selected autonomous recorder. We quantified vocal activity only during the before- and after-playback periods because temporal and spectral overlap between the broadcast stimulus and birds’ responses precluded reliable detection and attribution of vocalisations during playback, particularly for individuals vocalising farther from the loudspeaker. Thus, we quantified the number of songs produced by the target species during 10 min before and after playback. A song bout was defined as continuous vocalisation separated from the next bout by more than 3 seconds of silence. Song duration was measured in Raven Pro by placing a selection box around the complete song and recording the interval between the onset of its first element and the offset of its final element. All measurements were made by one observer (KB).

Video recordings from infrared camera traps were analysed by one observer (KB) using DaVinci Resolve 17 software. The dataset comprised 30 one-minute video clips for each experimental trial conducted at sedge warbler sites. Videos were viewed at up to 10 times their original playback speed to facilitate the initial detection of movement; each potential bird movement was subsequently verified at normal speed. For each clip, we recorded the number of flight events within the camera’s field of view. Species and individuals could not be reliably identified from the nocturnal video recordings because of limited image quality, infrared illumination, and the short duration of visible movement. Based on the silhouette and size of the bird observed in the video recordings, which are consistent with a sedge warbler, and on the fact that the experiment was conducted within the sedge warbler’s territory, we treated these detections as sedge warbler.

### Statistical analyses

Statistical analyses were restricted to the three species for which vocal activity was detected during the experimental periods: the sedge warbler, Savi’s warbler, and common snipe. The remaining seven species were excluded from species-specific models because no vocalisations were detected. For each analysed species, we modelled the number of songs recorded before and after playback periods, because vocalisations produced during the playback period could not be reliably distinguished from the broadcast stimulus. Moreover, because Savi’s warbler and sedge warbler songs have varying durations, we also built an additional model with the average song length as the response variable for these species. In this case, together with the songs-count model, this allowed us to test whether playback was associated with changes in song-bout number and duration. In all models, we used the same set of predictors, which were the experimental phase (2-level factor, before and after playback), night part (2-level factor, early night and late night), and sampling period (2-level factor, May and June). We also considered the interactions between experimental phase and night part, and experimental phase and sampling period because our hypotheses concerned whether responsiveness to playback varied over the course of the night and between sampling periods. All initial models included both hypothesised interactions. When models failed to converge or contained non-identifiable parameters, interactions were removed as necessary to obtain a reliable fit, while main effects involved in retained interactions were always kept. Experimental site identity was included as a random intercept to account for repeated observations collected at the same location. Because birds were not individually marked, this random effect represents repeated sampling of sites rather than confirmed repeated measurements of the same individuals. No common snipe vocalisations were detected during the June trials; consequently, the common snipe response model was restricted to the May sampling period.

Our experimental design allowed us to assess the response of sedge warbler flight activity to playback stimulation. We applied the same modelling framework as for the vocal activity models, using experimental phase, night part, sampling period, and their interactions as predictors, with experimental site identity included as a random intercept effect. Because no flights were recorded during the before-playback periods, this phase could not be included in the fitted count model. The final model therefore compared the during- and after-playback periods. The distribution of observed flight activity across all three experimental phases (before, during, and after playback) is presented separately using raw data to illustrate the full experimental context.

Models were fitted in R 4.5.2 (R Core Team, 2025) using the *glmmTMB* package (Brooks et al., 2017). Song- and flight-count models were analysed using negative binomial generalised linear mixed models (quadratic parameterisation; nbinom2) with a log-link function, whereas the average song duration response was analysed using a Gamma GLMM with a log-link function. Moreover, the sedge warbler song abundance model underpredicted zeros (45 observed vs. 25 predicted) and showed underdispersion (dispersion ratio = 0.246, p < 0.001). We therefore separated the response into two components: a binary model describing the probability of singing (binomial part) and a count model for the number of songs among observations with > 0 songs. The positive-count model remained strongly underdispersed (dispersion ratio = 0.464, p < 0.001). We therefore retained only the binary model for inference, as the available data did not support reliable modelling of song abundance among singing individuals due to extremely low variation. Model assumptions were evaluated using model summaries and dispersion and zero-inflation diagnostics using the *performance* package (Lüdecke et al., 2021). The significance of fixed effects was assessed using Type III Wald χ^2^ tests implemented in the *car* package (Fox & Weisberg, 2011), and estimated marginal means and pairwise comparisons were obtained using the *emmeans* package (Lenth & Piaskowski, 2017). The R project containing the raw data is hosted on GitHub at the link: https://github.com/kinkul1/Species-specific-responses-of-diurnal-birds-to-nocturnal-conspecific-song-playback.git

## RESULTS

Of the 10 tested bird species, only three species vocalised during the experiment, both before and after the playback: sedge warbler, Savi’s warbler, and common snipe. The other tested species did not sing even once during any experimental phase.

### Sedge warbler

The probability of singing in sedge warbler differed between sampling periods and between parts of the night (Fig. 1). Singing was less likely in June than in May (χ^2^ = 5.98, p = 0.014; Fig. 1), and more likely during late- than early-night trials (χ^2^ = 13.23, p < 0.001; Fig. 1). We found no clear evidence of an overall difference between before- and after-playback periods (χ^2^ = 2.98, p = 0.084; Fig. 1). Model estimates indicated high probabilities of singing in May in both phases (before: 0.93, after: 0.96), and lower in June (before: 0.73, after: 0.84). Pairwise comparisons showed no significant differences between phases within either month (May: z = −0.93, p = 0.35; June: z = −1.12, p = 0.26).

**Figure 1.**
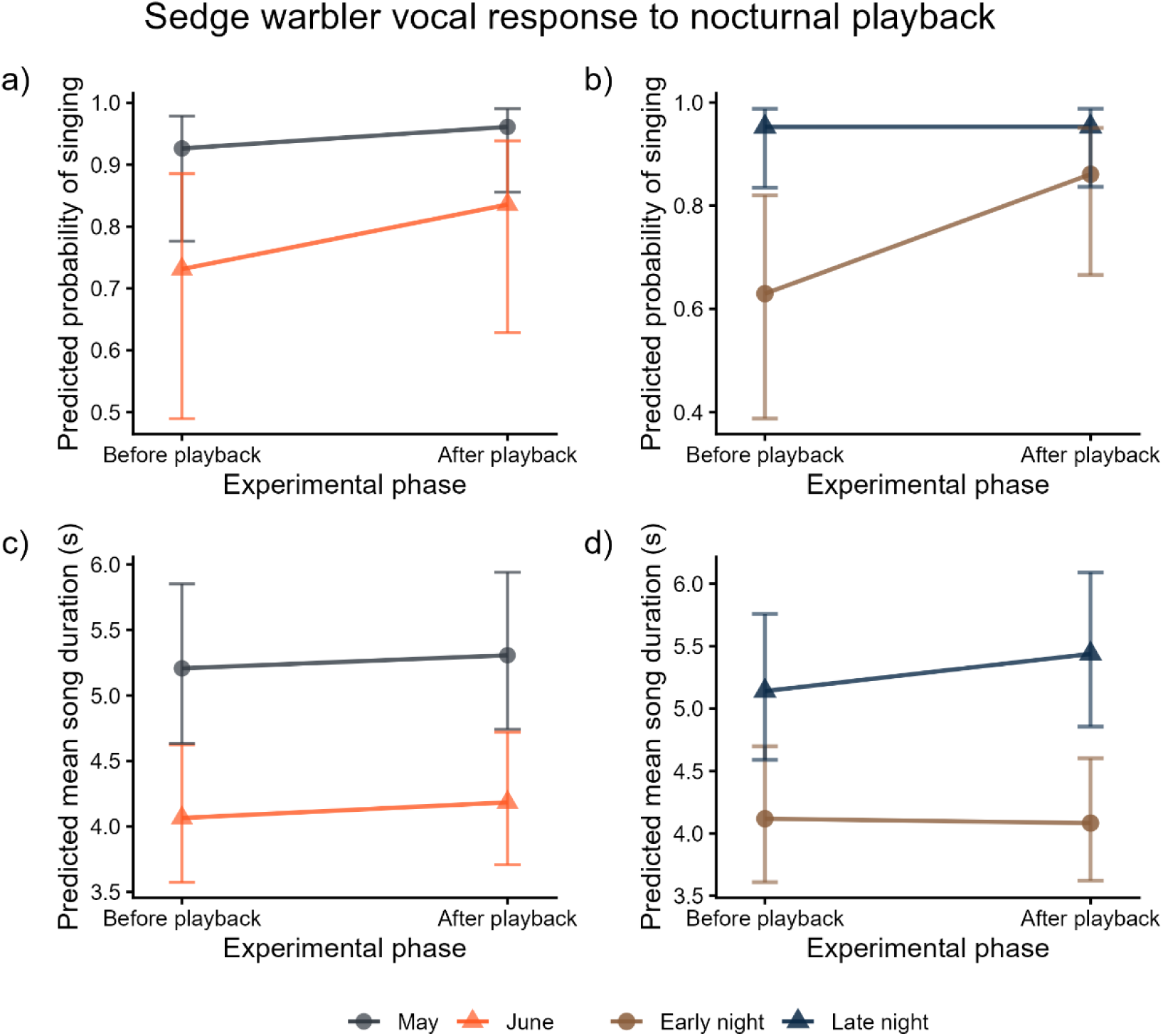
Predicted vocal responses of sedge warblers to nocturnal conspecific song playback. Model-estimated responses (±95% confidence intervals) showing changes in probability of singing (a–b) and mean song duration (c–d) before and after playback. Panels (a) and (c) show interactions between playback phase and sampling period, while panels (b) and (d) show interactions between playback phase and night part. Colours indicate months (May and June) or night periods (early and late night), respectively. Predictions were obtained from GLMMs that included experimental site identity as a random effect.

Average song duration did not differ between playback periods (χ^2^ = 0.002, p = 0.968). We also found no evidence that playback-phase effect varied between sampling periods (phase × sampling period: χ^2^ = 0.01, p = 0.92) or between parts of the night (phase × night part: χ^2^ = 0.44, p = 0.505). Moreover, estimated song-duration ratios showed no consistent change between the before- and after-playback periods in either May: after vs before, ratio = 1.02, z-ratio = 0.294, p = 0.768; June: after vs before, ratio = 1.03, z-ratio = 0.397, p = 0.691; Fig. 1c).

Raw observations showed that playback stimulation increased flight activity relative to the before-playback period, when no flights were recorded within the camera’s field of view (Fig. 2a). Because all before-playback observations were zero, the count GLMM was restricted to the playback and after-playback periods and therefore did not test the difference between the before-playback period and the subsequent phases. Flight activity increased during playback, followed by a decrease after playback in May, while in June it remained at a similarly elevated level after the experiment. Within the playback and after-playback phases, flight intensity was higher in June than in May (β = 0.87 ± 0.39 SE, z = 2.22, p = 0.027) and lower during late- than early-night trials (β = −0.77 ± 0.39 SE, z = −1.99, p = 0.047). We found no evidence that flight intensity differed between playback and after-playback phases (χ^2^ = 1.17, p = 0.279), or that the difference between these phases varied with sampling period (phase × sampling period: χ^2^ = 0.68, p = 0.41) or night part (phase × night part: χ^2^ = 0.60, p = 0.44). This pattern indicates that flight activity remained at a similar level after playback rather than decreasing immediately following the stimulus (Fig. 2b).

**Figure 2.**
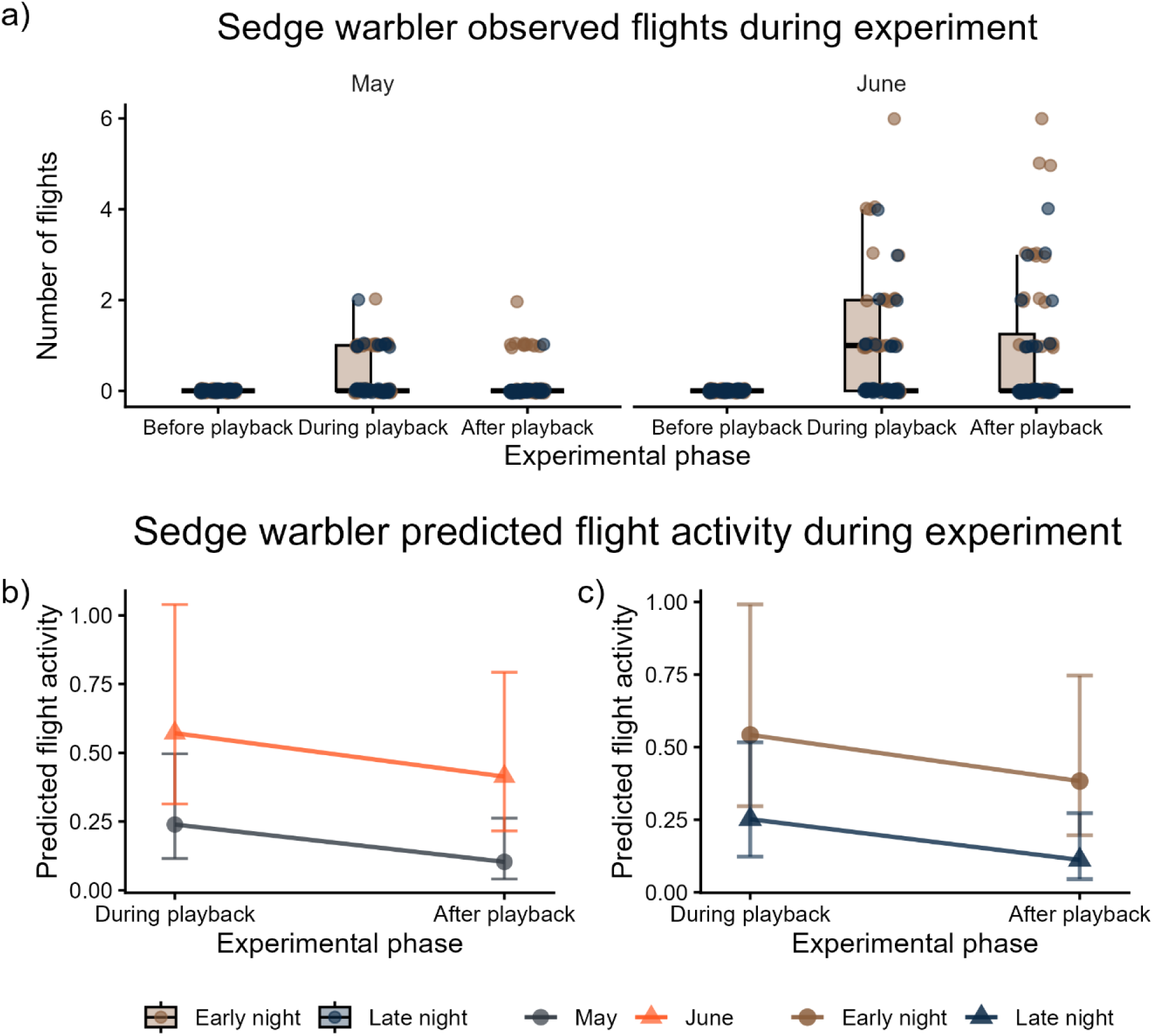
Flight activity recorded at sedge warbler sites during nocturnal conspecific song playback experiment. (a) Observed flight activity before, during, and after playback. Dots represent individual observations defined as the total number of flight events recorded within the camera’s field of view during each 10-min period. (b–c) Model-predicted flight activity during and after playback, shown by sampling period (May and June; b) and night part (early and late; c). Points represent predicted mean flight activity with 95% confidence intervals. The before-playback phase was not included in the GLMMs because no flights were recorded during this period (see Methods).

### Savi’s warbler

The number of songs differed between the before- and after-playback phases (χ^2^ = 4.26, p = 0.039; Fig. 3a) and was lower in June than in May (χ^2^ = 18.48, p < 0.001; Fig. 3a). Birds sang more after the playback than before in May (emmeans ratio before/after = 0.375, SE = 0.171, z-ratio = -2.146, p = 0.032; Fig. 3a), while no difference between experimental phases was observed in June (emmeans ratio before/after = 1.353, SE = 0.761, z-ratio = 0.538, p = 0.591; Fig. 3a). This pattern suggest that an after-playback increase was detected in May but not in June, possibly because overall singing intensity was low in June (Fig. 3). Consistent with this pattern, the experimental phase × month interaction showed marginal evidence of month-level heterogeneity (χ^2^ = 3.13, p = 0.077; Fig. 3b). Estimated marginal means showed that in May, the number of songs increased from 6.84 (before) to 18.21 (after), whereas in June, singing intensity remained low and similar between phases (0.61 vs 0.45). Night Part had no effect on song number (χ^2^ = 0.017, p = 0.897).

**Figure 3.**
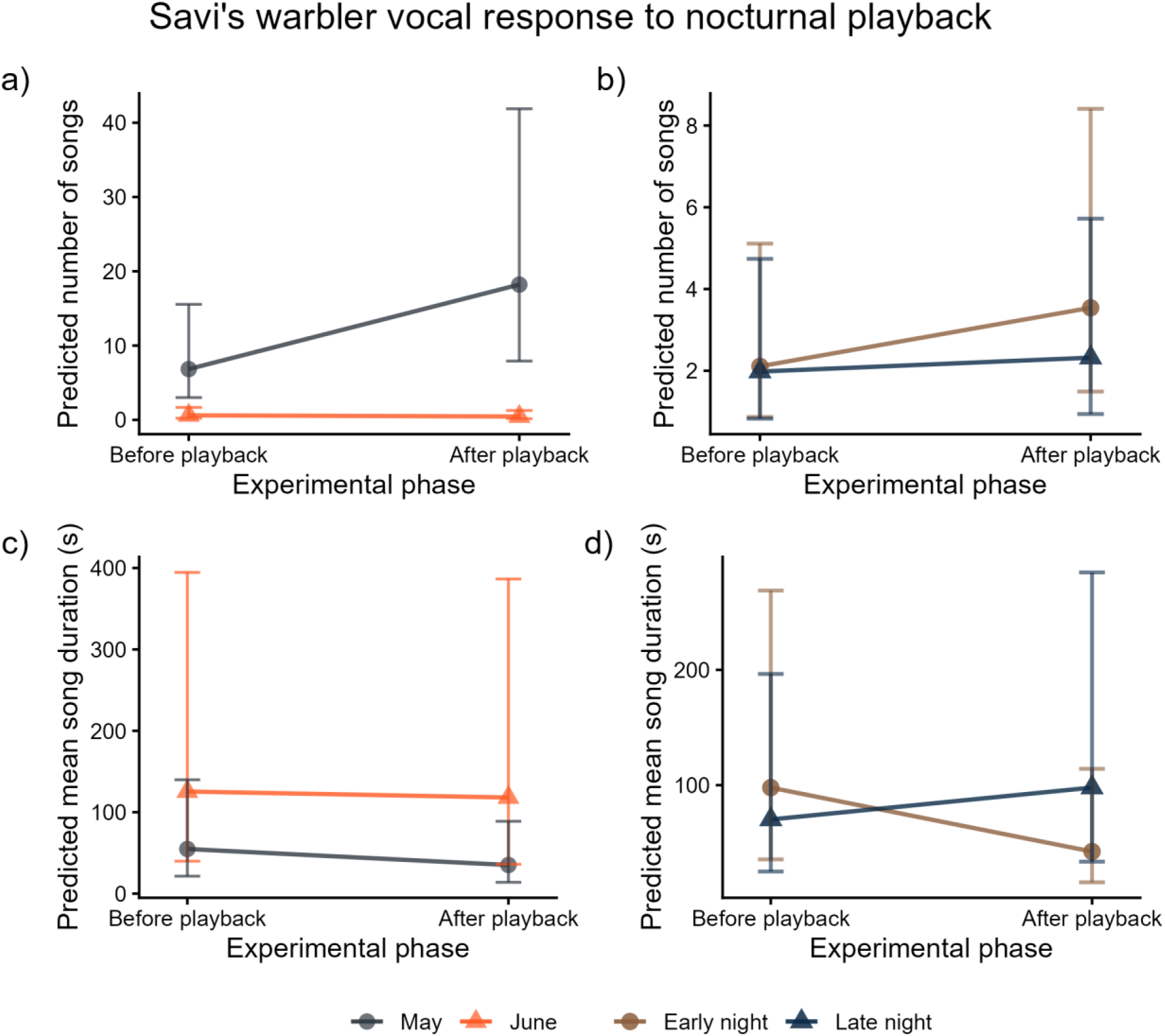
Predicted vocal responses of Savi’s warblers to nocturnal conspecific playback. Model-estimated intensity of singing (a–b) and mean song duration (c–d), with ±95% confidence intervals, during the before- and after-playback periods. Panels (a) and (c) show predictions by sampling period (May and June), whereas panels (b) and (d) show predictions by night part (early and late). Colours distinguish sampling periods in panels (a) and (c) and night parts in panels (b) and (d). Predictions were obtained from GLMMs that included experimental site identity as a random effect.

Among observation periods in which singing occurred, mean song duration differed between sampling periods, with shorter songs in May than in June (β = −1.211 ± 0.449 SE, z = −2.7, p = 0.007). There was no evidence of an overall difference in song duration between the before- and after-playback phase (χ^2^ = 0.11, p = 0.737), or that this difference varied between sampling periods (phase × sampling period: χ^2^ = 0.5, p = 0.479). However, the change in song duration between the before- and after-playback periods differed between the early and late parts of the night (phase × night part: χ^2^ = 6.24, p = 0.012; Fig. 3d). During early-night trials model-estimated mean song duration decreased from 97.9 s before playback to 42.4 s after playback (z-ratio = −2.4, p = 0.016; Fig. 3d). During late-night trials the estimated mean increased from 70.2 s before playback to 97.9 s after playback, but this contrast was not significant (z-ratio = 0.896, p = 0.37; Fig. 3d).

### Common snipe

Because no common snipe vocalisations were detected in June, the analysis was restricted to the May sampling period. The initially specified phase × night-part interaction could not be estimated reliably because sparse observations for certain factor combinations resulted in high collinearity (high VIF values) among model coefficients. The model-estimated number of vocalisations increased from 0.14 during the before-playback period to 4.22 during the after-playback period (χ^2^ = 4.34, p = 0.037; Fig. 4). The number of vocalisations did not differ between the early and late parts of the night (χ^2^ = 0.40, p = 0.526). Thus, common snipe vocal output was higher after playback than before playback during the May trials, whereas no association with night part was detected. Model diagnostics indicated no evidence of overdispersion (dispersion ratio = 0.155, p = 0.896) or zero inflation (observed/predicted zero ratio = 1.00).

**Figure 4.**
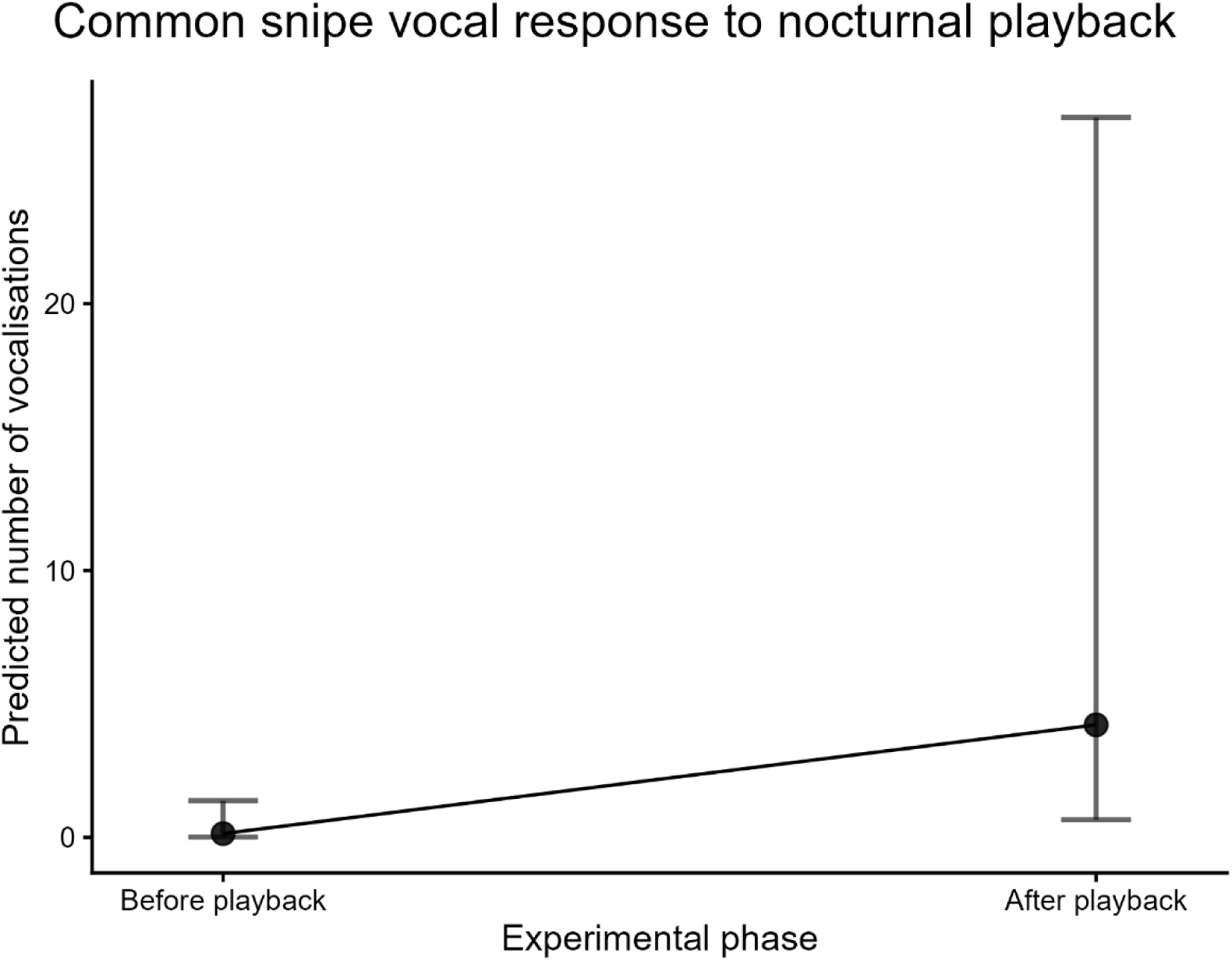
Predicted vocal response of common snipe to nocturnal conspecific playback. Model-estimated vocalisation intensity (±95% confidence intervals) before and after playback. Predictions were obtained from a GLMM that included experimental site identity as a random effect.

## DISCUSSION

We recorded nocturnal vocal activity in three of the 10 species included in the nocturnal playback experiment: the sedge warbler, Savi’s warbler, and common snipe. The occurrence and form of responsiveness differed among these species. Savi’s warblers produced more songs after than before playback during the May sampling period, and common snipes showed a corresponding after playback increase in May, the only sampling period in which this species vocalised. By contrast, sedge warblers showed no clear change in the probability of singing or in mean song duration following playback. These results indicate that some otherwise diurnal species perceive conspecific song broadcast at night as behaviourally relevant, but they provide no evidence for a uniform response across species.

In the case of the sedge warbler, we observed a tendency to sing with higher probability after than before playback stimulation, but the phase effect was statistically insignificant, and pairwise comparisons did not identify before to after playback difference in either sampling period (Fig. 1). Thus, the vocal data do not provide evidence that sedge warblers increased their probability of singing or song duration following playback. No flight events were detected before playback at sedge warbler sites, while their activity suddenly increased during the playback and slowly decreased after the playback (Fig. 2). Flight counts recorded during and after playback showed the opposite temporal association: within these two phases, fewer events were detected during late- than early-night trials. This contrast could reflect temporal changes in response mode, with birds relying relatively more on movement early in the night and vocal activity towards dawn. In our experiment, species identity could not be determined reliably from the infrared recordings, however based on loudspeaker located within the sedge warbler territory, size and silhouette of flying birds, we assumed that it was highly likely that flying birds are sedge warbler. Daytime playback experiments in sedge warblers have shown that conspecific song can elicit strong aggressive responses from territorial males, although the strength of this response declines after pairing (Catchpole, 1977). Another study confirms that sedge warbler diurnal songs are also directed at rival males and serve the dual functions of mate attraction and territory defence (Brumm et al., 2011). Studies on the ovenbird *(Seiurus aurocapilla*) have shown that birds do not respond aggressively to the night song of foreign males, suggesting that nocturnal singing in this species is used for mate attraction rather than territory defence (Foote et al., 2018). Importantly, those findings do not by themselves demonstrate mate attraction; they principally indicate a lack of support for the tested intraspecific response. In field sparrows (*Spizella pusilla*), males did not countersing to octurnal playback, while movement responses—particularly those of females—varied with breeding stage, making an intersexual interpretation more plausible than a conventional territorial one. In the common nightingale (*Luscinia megarhynchos*), unpaired males sing intensely at night (Amrhein et al., 2002). This behaviour becomes intermittent after pair formation but resumes when a male loses his mate. By contrast, territory-prospecting males are more active around dawn, when both paired and unpaired territory holders sing at high rates. This correspondence between receiver activity and male signalling suggests that nocturnal and dawn song may serve different dominant functions even within the same species. Thus, the observed pattern—in which sedge warblers did not increase vocal output, whereas some individuals appeared near the speaker, may suggest that nocturnal song in sedge warblers is directed predominantly towards females. This hypothesis needs further examination.

The increases in vocal output after playback by Savi’s warbler and common snipes are consistent with a social response to conspecific song (Fig 3., Fig 4.). In Savi’s warblers, mean song duration also decreased after playback during early-night trials, whereas no corresponding decrease was detected during late-night trials. In some bird species, shorter or structurally simpler songs are associated with close-range male–male interactions and aggressive contexts. For example, playback studies of Acrocephalus warblers have shown that males may produce shorter, simpler song types during territorial encounters (Catchpole & Leisler, 1989). Another study showed that polygynous species of great reed warbler (*Acrocephalus arundinaceus*) has evolved short, simple and stereotyped songs for territorial defence through intrasexual selection (Catchpole, 1983). Thus, vocal responses of Savi’s warbler and common snipes indicate that playback was followed by increase vocal output, a pattern consistent with a territory defence function. Therefore, diversity of observed behavioural patterns suggests that the functions of nocturnal singing in diurnal birds are species-specific, influenced by multiple factors and may contribute to both mating-related communication and territorial defence, like in willie wagtails (*Rhipidura leucophrys*)(Dickerson et al., 2023).

Nocturnal vocal activity also varied over the course of the night. Sedge warblers were more likely to sing during late- than early-night trials, independent of playback (Fig. 1). A similar increase towards dawn has been reported in broader monitoring of nocturnal singing in Central Europe and may be associated with increasing natural illumination or the gradual transition into the dawn chorus (Buda et al., 2025). In contrast, in Savi’s warbler and common snipe we found no differences in vocalization intensity between the early- and late-night periods.

Furthermore, the function of nocturnal singing may change across the breeding season. Sedge warblers were more likely to sing in May than June, Savi’s warblers produced more songs in May, but they were shorter than in June, and common snipe vocalisations were observed only in May. Across all these three species examined, nocturnal vocalisation was more frequent at the beginning of the breeding season. Greater nocturnal activity early in the season could be associated with territory establishment, mate acquisition, a greater prevalence of unpaired individuals or egg-laying. Reproductive context can affect nocturnal activity in other diurnal species, but the direction and interpretation of these relationships vary. Yellow-breasted chats (*Icteria virens*) undertake nocturnal extraterritorial movements, and female activity is concentrated particularly during the fertile period, which is consistent with nocturnal mate prospecting or extrapair mating opportunities (Ward et al., 2014). These findings concern movement and mating context, rather than demonstrating that males increase night song specifically to prevent extrapair copulations. Similarly, male willie wagtails increased nocturnal singing during the fertile periods of resident females, a pattern consistent with several non-exclusive functions, including mate stimulation, mate guarding, and post-pairing attraction. These examples reinforce the need to measure mating status, female fertility, and receiver behaviour directly before assigning a reproductive function to nocturnal song.

Despite several limitations, including the lack of individual marking, the inability to identify reliably the species or sex of birds recorded by the infrared cameras, and the absence of information on breeding status, our study shows that diurnal birds modify their behaviour in responses following playback of an unfamiliar conspecific male’s song at night. Savi’s warblers and common snipes produced more vocalisations after playback during the May sampling period, whereas sedge warbler vocal responses were not clearly affected, but birds approach to the loudspeaker during and after playback stimulation. Nocturnal activity also varied with sampling period and, for sedge warblers, with night part. These findings support the view that nocturnal vocal communication is context- and species-dependent, consistent with recent experimental and comparative work showing substantial variation in nocturnal responsiveness, mating context, environmental correlates, and potential costs. The overall results suggest one of the major roles of singing at night for these species is to set and defend territory, as, besides playback response, the overall singing activity was higher at the start of the breeding season when establishment and maintenance of the pair bond occur. Additionally, the lack of vocal response to playback in the sedge warbler may indicate that, in this species, nocturnal singing serves to attract females. These findings indicate that nocturnal singing may represent an important behavioural adaptation that may have context-dependent social functions that should be tested against direct measures of mating, territorial behaviour and fitness.

## Acknowledgements

A study financed by the Ministry of Education and Science in Poland budget funds for science in 2022-2024 as a research project under the “Diamond Grant” program (0128/DIAa/2019/48).

